# Cooperative changes in solvent exposure identify cryptic pockets, conformational switches, and allosteric coupling

**DOI:** 10.1101/323568

**Authors:** Justin R. Porter, Katelyn E. Moeder, Carrie A. Sibbald, Maxwell I. Zimmerman, Kathryn M. Hart, Michael J. Greenberg, Gregory R. Bowman

## Abstract

Proteins are dynamic molecules that undergo conformational changes to a broad spectrum of different excited states. Unfortunately, the small populations of these states make it difficult to determine their structures or functional implications. Computer simulations are an increasingly powerful means to identify and characterize functionally-relevant excited states. However, this advance has uncovered a further challenge: it can be extremely difficult to identify the most salient features of large simulation datasets. We reasoned that many functionally-relevant conformational changes are likely to involve large, cooperative changes to the surfaces that are available to interact with potential binding partners. To examine this hypothesis, we introduce a method that returns a prioritized list of potentially functional conformational changes by segmenting protein structures into clusters of residues that undergo cooperative changes in their solvent exposure, along with the hierarchy of interactions between these groups. We term these groups exposons to distinguish them from other types of clusters that arise in this analysis and others. We demonstrate, using three different model systems, that this method identifies experimentally-validated and functionally-relevant conformational changes, including conformational switches, allosteric coupling, and cryptic pockets. Our results suggest that key functional sites are hubs in the network of exposons. As a further test of the predictive power of this approach, we apply it to discover cryptic allosteric sites in two different β-lactamase enzymes that are widespread sources of antibiotic resistance. Experimental tests confirm our predictions for both systems. Importantly, we provide the first evidence for a cryptic allosteric site in CTX-M-9 β-lactamase. Experimentally testing this prediction did not require any mutations, and revealed that this site exerts the most potent allosteric control over activity of any pockets found in β-lactamases to date. Discovery of a similar pocket that was previously overlooked in the well-studied TEM-1 β-lactamase demonstrates the utility of exposons.

## Introduction

Proteins are highly dynamic molecules that are capable of accessing a wide variety of excited conformations. Many of these excited states have important biological functions. For example, many proteins predominantly adopt an off state until they interact with a binding partner that stabilizes a higher energy on state. However, the most common tools for structural biology, such as x-ray crystallography and cryoelectron micrography, typically only provide a static picture of one (or a few) low energy states.

Computer simulations, because of their excellent spatiotemporal resolution, are a promising means to identify functionally-relevant excited states and conformational changes (1). However, simulations have historically faced severe limitations. In particular, the inability to capture slow processes, such as large-scale conformational changes, has hampered the routine discovery of physiologically-relevant excited states using computer simulations. However, enormous advances in computer hardware and simulation algorithms have made it possible to capture processes that occur on tens to hundreds of milliseconds, finally giving access to this physiologically-important timescale for many proteins (2). One successful approach has been to combine many parallel simulations executed on commodity hardware into a single model of protein dynamics using Markov state models (MSMs) (3). MSMs are network models of protein energy landscapes composed of many conformational states and the probabilities of hopping between them. Because they are able to integrate information from many independent simulations, they are capable of reaching timescales many orders of magnitude larger than any of the individual simulations used to build the model (1, 3-6).

The growing availability of long-timescale simulations has revealed a new major challenge: extracting meaningful insights from the resulting colossal datasets. These datasets are not only composed of hundreds of millions or billions of timepoints, but are also embedded in tens or hundreds of thousands of dimensions. Numerous methods have been developed to address this challenge. One method, projecting simulation data onto specific order parameters is a valuable means to test hypotheses, but this approach requires detailed foreknowledge of which parameters are important to avoid obscuring important features (7). For data sets where foreknowledge is not available, unsupervised methods have been developed to learn what degrees of freedom are important. For example, principle component analysis (PCA) (8) highlights large geometric changes. Unfortunately, larger conformational changes are not necessarily more functionally relevant. For example, the large variance in the atomic positions of a disordered loop can easily dwarf a more subtle, but more functionally relevant, conformational change. Another approach, time-lagged independent component analysis (tICA) (9, 10) instead focuses on slowly varying dimensions. Slowness, however, does not necessarily imply functional relevance. For instance, the process of flipping a phenylalanine about its ring axis may be slow, but the exchange-symmetry of atoms on either side of the ring means the process does not alter the conformation at all.

To address these challenges, we hypothesized that functionally-relevant conformational changes are likely to result in large, cooperative changes to the surfaces of a protein that are available to interact with potential binding partners. This hypothesis was inspired, in part, by the fact that surface chemistry is an especially important feature of most proteins, since it is how the protein interacts with other objects, including any substrates and binding partners. Furthermore, we reasoned that functionless cooperativity at protein surfaces is rare. Physically, the folded state is bombarded by thermal noise which—in the absence of a specific design constraint—will tend to decorrelate any arbitrary pair of features. Genetically, sequence drift would be expected to eliminate cooperativity that is not selected for over time; much as early protein redesigns sometimes inadvertently destroyed cooperative folding (11). This assumption is also the basis for sequence-based methods that use patterns of conservation and covariance to infer pairs of residues that are in direct contact in a protein’s three-dimensional structure, or that are allosterically coupled (12-14).

To make testing this hypothesis tractable, we developed a method that returns a prioritized list of potentially functional conformational changes by segmenting protein sequences into clusters of residues that undergo cooperative changes in their solvent exposure and uncovers the hierarchy of interactions between these groups. We term these clusters of mutually-correlating residues *exposons* to disambiguate them from other forms of clustering found in this work and in the literature. To identify exposons and the structural motions that give rise to them, we present an efficient, MSM-based approach.

For a concrete example of the utility of identifying cooperative changes in solvent exposure, consider cryptic pockets. Cryptic pockets are transient concavities on protein surfaces that open when the protein fluctuates to an excited state (15, 16). Pocket opening concomitantly increases the solvent exposure of surrounding residues, and closing a pocket simultaneously reduces their exposure. Thus, these residues undergo correlated changes in their solvent exposure and are likely to form an exposon.

To establish the value of exposons, we demonstrate that they naturally identify a variety of functionally-relevant conformational changes without any foreknowledge of what structural features are important for a given system. First, we show that exposons identify cryptic pockets and allostery in the enzyme TEM-1 β-lactamase. We then show that they detect a conformational switch in the Ebola virus’ nucleoprotein (eNP) and allostery in the catabolite activator protein (CAP). Then we use exposons to prospectively discover cryptic allosteric sites in two different β-lactamase enzymes with less than 40% sequence identity, TEM-1 and CTX-M-9, and perform *in vitro* biochemical experiments to test our predictions.

## Methods

### Simulations

As described previously (17), simulations were run at 300 K with the GROMACS software package (18) using the Amber03 force field (19) and TIP3P (20) explicit solvent. β-Lactamase simulations were deployed on the Folding@home distributed computing platform (21), while simulations of eNP and CAP were performed on NVIDIA P100 GPUs on our local cluster. For our retrospective work, we used previously published datasets including 90.5 μs of simulation of TEM-1 β-lactamase with the M182T substitution (17), 28.0 μs of simulation of eNP (22), and 1.5 μs of simulation of CAP (23). For our prospective work on CTX-M-9 β-lactamase, we ran 76.0 μs of aggregate simulation on Folding@home.

### Solvent exposure featurization

To generate a solvent exposure featurization of each dataset, we computed the solvent-accessible surface area (SASA) of each residue’s sidechain in each simulation frame to a drug fragment-sized probe using the Shrake-Rupley (24) algorithm, as implemented in MDTraj (25). The result is a set of *t* vectors of length *n*, where *n* is the number of residues and *t* is the number of frames. A probe size of 2.8 Å was chosen because previous work suggests this value identifies pockets that can accommodate a drug-sized molecule (15). Jug (26) was used to organize the parallel execution of many independent tasks, including parallelizing solvent-accessibility calculations across many cores.

### Markov state models

We defined MSM microstates by clustering the sidechain solvent-accessible surface area featurized representation. Using this featurized space, rather than a geometrical criterion like RMSD, to do our state space discretization synergizes with the calculation of exposons, described below. Clustering was performed with a hybrid *k*-centers/*k*-medoids algorithm (27) with a Euclidean distance metric. Five rounds of *k*-medoids updates were used, where updates were accepted if the largest distance to the nearest medoid decreased. Transition probabilities were estimated by row-normalizing the transition counts matrix after the addition of a pseudocount inversely proportional to the number of states (e.g. 1/*n*), as suggested in (28) and (29). The lag time and clustering stopping condition of 2.6 nm^2^ and .1 ns (TEM-1), 3.5 nm^2^ and 0.1 ns (CTX-M-9), 5.0 nm^2^ and 0.5 ns (eNP), and 3.7 nm^2^ and 0.4 ns (CAP) were chosen by the implied timescales test (Fig. S1). The highest flux pathways between two sets of states were then extracted using transition path theory (30, 31).

### Exposon calculation

Beginning with the featurized representation for the representative conformation for each MSM state, we classified each sidechain in each state as exposed or buried using a fixed threshold. We chose a fixed threshold rather than a continuous threshold to reduce the number of parameters (a sigmoid, for example, would require a step width and a step midpoint) and because previous work suggested that mutual information performs better when a smaller number of bins are used (23). Our fixed threshold in this paper was 2.0 Å^2^, but choices in the range 2.0–5.0 Å^2^, as well as values formulated as a fraction of maximum possible sidechain exposure in the 1-3% range, gave similar results. This invariance implies that our algorithm does not erroneously favor larger residues due to their larger maximum possible SASA. The end result is a featurization of each MSM state, wherein each snapshot is represented by a binary vector with one entry per residue that contains a one for exposed residues and a zero for buried residues.

We then calculate the mutual information between each pair of residues. Mutual information (MI) is a measure of the statistical interdependence of two random variables. It is given by the equation,

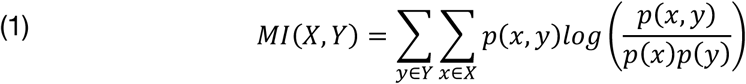

where *X* and *Y* are any pair of residues and *x* and *y* represent the solvent accessibility states (i.e. buried, exposed) of the corresponding residue. The probability *p*(*x*) is the probability that a residue is observed in state *x* and *p*(*x, y*) is the joint probability of *x* and *y*. These probabilities are the equilibrium probabilities calculated during MSM fitting. In computing mutual information in CAP, which is a symmetric homodimer, symmetry was enforced by reparametrizing *Y* in terms of whether or not it is on the same chain or the opposite chain as *X*.

Exposons are the cluster assignments computed by affinity propagation (32). We use the affinity propagation implemented in scikit-learn 0.19.0 (33) and zero initial affinities. The damping parameters were 0.9 for TEM-1, CTX-M-9 and eNP and 0.95 for CAP, generally chosen to be as high as possible (creating a low number of exposons) without causing the algorithm to converge on a single exposon for the entire protein. This typically generates 10-50 exposons, of which we visualize the top 3-10. Affinity propagation is robust to the choice of damping parameter, giving similar values for much of the range of valid choices (Fig. S2).

Coarse-grained exposon graphs are network models of the communication between each pair of exposons. In this model, each node represents an exposon and each edge represents the communication between a pair of exposons. The weight of each edge in the coarse-grained network is calculated by,

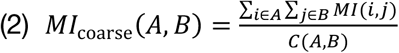

where *A* and *B* are any pair of exposons (sets of residues) and *C* is the channel capacity, which is the maximum possible mutual information between two pairs of exposons (23). In this case, the channel capacity is given by *C*(*A, B*) = min (|*A*|, |*B*|) or one bit per residue up to the number of residues in the smallest exposon. When visualizing coarse grain exposons, we omit self-edges and very low-valued edges (< 0.015 for TEM, < 0.075 bits for eNP, and < 0.2 bits for CAP).

Eigenvector centrality calculations were performed with NetworkX (34).

### Synthetic labeling

We predict time-dependent labeling behavior using the MSM we fit as described above. Synthetic labeling experiments are performed first by identifying all states in which the residue of interest is exposed and converting them to sink states by zeroing out the rows in the transition probability matrix. Then, iteratively multiplying the equilibrium probability distribution by this new matrix gives a monotonically decaying fraction of ‘unlabeled’ probability density as density flows into the sink states. Finally, we fit the unlabeled fraction as a function of time to a single exponential to yield a rate. In the limit of a perfectly good fit, this rate is equivalent to a mean first passage time (35). An implementation of this simple procedure is provided in a Jupyter notebook (see ‘Code Availability’). We used SciPy (36) version 0.19.1 for curve fitting.

### Protein expression and purification

TEM-1 was purified from the periplasmic fraction of BL21(DE3) cells (Agilent Technologies) using both cation exchange and size exclusion chromatography. The full protocol is described in previous work (37).

We subcloned the CTX-M-9 gene into the multiple cloning site of pET9-a vector. Plasmids were transformed into BL21(DE3) Gold cells (Agilent Technologies) for expression under T7 promoter control. Cells were induced with 1mM IPTG at OD=0.6 and grown for 5 hours at 37°C. The cells were then centrifuged and the pellet was frozen at −80 °C.

CTX-M-9 cells were resuspended in 20 mM sodium acetate, pH 5.5, sonicated, and then centrifuged. The protein was purified from the insoluble cytoplasmic fraction. The pellet was unfolded in 9 M urea 20 mM sodium acetate, pH 5.5 and centrifuged. CTX-M-9 was then refolded in 20 mM sodium acetate, pH 5.5, purified by both cation exchange and size exclusion chromatography and stored similarly to TEM-1.

### Thiol labeling

We observe the change in absorbance over time of DTNB (Ellman’s reagent, Thermo Scientific), a small molecule that changes its absorbance as it covalently binds reduced cysteine sidechains (15). We used a SX20 stopped-flow instrument (Applied Photophysics) with a dead time of 1.5 ms. Measurements were taken over time in 20 mM Tris, pH 8 1% DMSO, followed at an absorbance of 412 nm (ε_412_ = 14,150 M^-1^ cm^-1^), and fit by a single exponential (Fig. S3). Our previous work with thiol labeling was performed using manual mixing in a standard UV-Vis spectrophotometer (15), but in this work we used a stopped-flow instrument that gives access to faster timescale motions and improves the quality of fits because it offers a dead time that is much shorter than the time scale of our experiments. It also allows for the use of lower DTNB and protein concentrations.

The labeling rate at a given DTNB concentration can be described by the Linderstrøm-Lang model, originally derived for hydrogen-deuterium exchange (38):

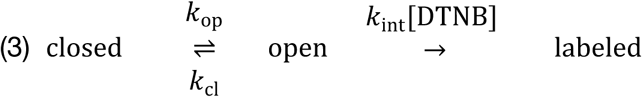

The observed rate is given by:

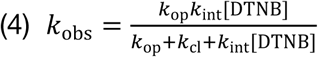

which is a nonlinear function that approaches a linear dependence on [DTNB] at low concentrations and [DTNB] independence at high concentrations. In the limiting case where *k*_cl_ ≪ *k*_int_[DTNB] called the EX1 regime, the observed rate of labeling reduces to

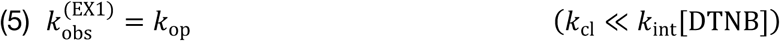

In the limiting case where *k*_cl_ ≫ *k*_int_[DTNB] called the EX2 regime, the observed rate of labeling reduces to

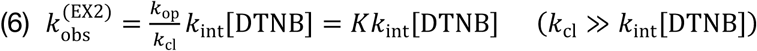

where *K* is the equilibrium constant between the open and closed forms. In the intermediate regime where *k*_cl_ ≈ *k*_int_[DTNB], called the EXX regime, one must fit to the full expression (given in equation 3). We found that over the concentrations of DTNB used, TEM-1 S243C labeling was in the EX2 regime (linear dependence on [DTNB]) and CTX-M-9 labeling was in the EXX regime (nonlinear dependence on [DTNB])

The three regimes differ in terms of the controls required to demonstrate that labeling is not occurring predominantly in the unfolded state. In the EX1 regime, the observed labeling rate for a pocket must be faster than the rate of global unfolding. Neither of the pockets we test in this paper labeled in this regime, but we have previously observed labeling rates in pockets that showed this behavior (15). In the EX2 regime, the equilibrium constant for pocket opening must be greater than that for the unfolded state (Equation 5). To determine these quantities for TEM-1 S243C, we measured the *K* of unfolding (Table S1) and the intrinsic rate of labeling for the denatured protein. To determine the intrinsic rate of labeling, *k*_int_, our labeling assay was repeated with the addition of 6 M urea. In the EXX regime, the observed labeling rate for a pocket must be greater than the maximum expected labeling rates of the unfolded state in either the EX1 or EX2 regimes (derivation in SI). Thus, to test that CTX-M-9’s labeling rate is not consistent with global unfolding alone, we measured both its rate of unfolding (Fig. S4) and its thermodynamic stability (Table S2). We then combined that with the fit value of *k*_int_ (Table S1) to produce a piecewise function that is an upper bound for labeling from the unfolded state. In this case, however, we found that the unfolding rate is the relevant control for all DTNB concentrations used here—the population of unfolded enzyme is relevant only at DTNB concentrations less than about 170 nM, the DTNB concentration where *k*_cl_ = *k*_int_.

### Urea melts and unfolding kinetics

Equilibrium stabilities and unfolding kinetics were acquired on a Chirascan circular dichroism spectrometer (Applied Photophysics) at a temperature of 25°C. Protein denaturation was observed by measuring the average ellipticity over 60 seconds at 222 nm as a function of urea concentration (Fig. S5, Table S2). Samples of 35 μg/ml protein were equilibrated in 50 mM potassium phosphate pH 7 and varying concentrations of urea overnight prior to data collection.

To determine the global unfolding rate, we used a linear extrapolation model (39) fit the log observed unfolding rates as a function of urea concentration at concentrations above the concentration at which it is half folded and half unfolded (the C_m_, Table S2) and extrapolated back to 0 M urea (Fig. S4). Concentrations were between 4 and 5.5 M urea for TEM-1 M182T and between 1.8 and 2.8 M urea for CTX-M-9.

### Activity measurements

Activity measurements were performed on both labeled and unlabeled protein. In order to measure the activities of the labeled proteins, 10 μM S243C and 5 μM CTX-M-9 were each incubated with excess DTNB for one hour, giving ample time for both proteins to fully label prior to the activity measurements. The proteins were then separated from excess DTNB using size exclusion chromatography.

Enzyme activities against nitrocefin (Cayman Chemical Company) were monitored at 482 nm (ε_482_ = 15,000 M^-1^ cm^-1^) using a Cary 100 UV-vis spectrophotometer (Agilent Technologies). Reactions were measured in 50 mM potassium phosphate, 10% glycerol (v:v), 2% DMSO pH 7.0 at 25 °C using 2 nM enzyme. Initial velocities were plotted as a function of nitrocefin concentration and fit to a Michaelis–Menten model to extract k_cat_ and K_m_ values (Fig. S6, Table S3).

### Visualizations

Protein structures were visualized using PyMOL 2.2 (40). Graphs were embedded with Fruchterman-Reingold algorithm (41) in NetworkX (34).

### Code availability

Code is available on GitHub as `bowman-lab/enspara`. The analysis described in this manuscript is reproduced in miniature, complete with example parameter choices and visualization recommendations, in a Jupyter notebook in the `enspara/examples` directory.

## Results

### Exposons simultaneously capture conformational changes and allosteric communication at protein surfaces

To identify exposons, we first construct an MSM (see *Methods*). In this work, we defined states using the Euclidean distance between vectors of sidechain solvent accessible surface areas, but in general, other methods can also be used. Each state in the MSM is then represented by a binary vector that characterizes the solvent exposure of each residue in the cluster center for that state (i.e. element *𝑖* is zero if residue *𝑖* is buried or one if it is exposed) (Fig. 1A). Based on previous work (15), sidechains are classified as exposed if their surface area exceeds 2 Å^2^. We then compute the mutual information (MI) between each pair of residues, giving a square MI matrix (Fig. 1B). Mutual information—defined in Eq 1 of *Methods*—is a nonlinear measure of the statistical interdependence of two random variables that has been previously used in studies of protein allostery (23, 42). A particularly useful property of the mutual information is that residues that never change their solvent exposure (i.e. have zero entropy) have zero mutual information. Finally, we cluster this mutual information matrix using affinity propagation (32) to assign residues to exposons. The resulting mutual information matrix can be visualized as a network with nodes colored according to their exposon assignment (Fig. 1C). The list of exposons can be prioritized for further analysis based on the total information of each exposon, which is the sum of mutual information between each residue in that exposon and any other exposon (23). This tends to identify larger exposons with more communication, which we reasoned are more likely to be functionally relevant and less susceptible to noise and errors introduced by finite sampling.

**Figure 1:**
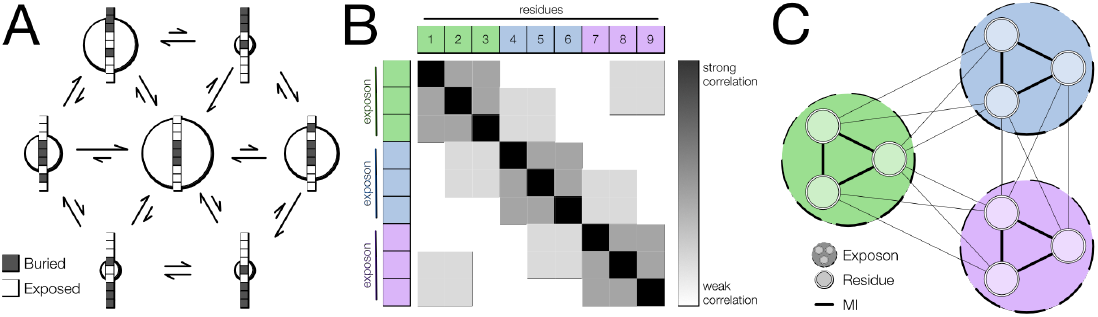
A schematic outline of our method for identifying exposons. (A) A Markov state model composed of variably populated states (circles, population is indicated by circle diameter) and transitions between states (single-headed arrows, probability is indicated by arrow length). Each state is associated with a binary exposed/buried classification for each residue, indicating whether the residue is exposed or buried in that state (column of black and white boxes, white denoting buried and black denoting exposed). (B) An all-against-all pairwise mutual information (MI) matrix that is calculated from (A). Exposons, indicated by the colored groups in the margins, are clusters of residues with mutually high pairwise mutual information. (C) The residue-level network representation of a mutual information matrix. Residues are indicated by double-edge circles, and are colored by their exposon membership. Exposons are indicated by dashed-lined circles. The mutual information between residues is indicated by straight lines between double-edged circles, and the weight of the line represents the strength of the correlation.

Once exposons have been computed, we typically wish to identify—in terms of atomic coordinates and protein conformations—which motion or motions give rise to an exposon. MSMs, which contain kinetic as well as thermodynamic information, are a natural source of this information. An MSM’s top eigenmodes capture how much each conformational state participates in the slowest motions observed in a simulation. Specifically, the state’s with the lowest eigenvector components are interconverting with the states with the highest eigenvector components. Therefore, we reasoned that an MSM’s top eigenmodes provide a facile means to identify the dominant motions contributing to an exposon (3). To identify which eigenmode reports on a particular exposon, we first compute the degree to which changes in an exposon’s solvent exposure are correlated with each eigenmotion in an approach similar to dynamical fingerprinting (43). We then choose the eigenmode that maximizes this correlation and extract the structures of the conformers at the extremes of this motion.

This conceptual framework has several important advantages over more traditional geometric approaches. First, it does not make any assumptions about which types of surfaces are most interesting—instead, any surfacial rearrangement that shows cooperativity will be detected. Second, this approach explicitly considers the entire sampled ensemble and uses this information to prioritize the most interesting features *of the ensemble*, rather than relying on structural features of particular conformers. Third, because exposons exist in sequence space, the results are insensitive to structural alignments and can be easily compared with experimental techniques that provide a read-out at the primary structural level, including thiol labeling. Consequently, this tool is applicable to a wide variety of conformational changes and scientific questions, as we demonstrate below.

Our approach is predicated on the assumption that interesting features are those that change at the surface. Our motivation for this assumption was that most interesting protein behavior ultimately is a consequence of the protein’s ability to interact with other objects, which occurs at the surface. Any cooperative rearrangement that does not substantially alter a protein’s solvent exposure will not be detected. For instance, a rotameric transition that creates geometry necessary for catalysis may not entail any change in solvent exposure. Likewise, any cooperative changes occurring exclusively in the protein core will not be detected. Allosteric coupling between two surface sites that occurs through the core will be detected, but the mechanism will not be apparent since exposons will only be sensitive to the end-points. Another potential limitation of our approach is imposed by the use of mutual information, which is only sensitive to features that change. A concavity at a protein surface that never changes its conformation will not be identified by this method—this situation is much better suited to the many excellent geometrical pocket detection methods proposed over the years (44-47).

### Retrodiction of a cryptic allosteric site in TEM-1 β-lactamase

As a first test of our model, we examined its ability to identify cryptic allosteric sites. A cryptic allosteric site is a pocket that is absent in available structures but is present in excited states and can exert allosteric control over a distant functional site, such as an enzyme’s active site. Cryptic pockets are a particularly interesting class of excited states because identifying new cryptic sites could offer new druggable sites on established drug targets, provide a means to inhibit targets that are currently considered undruggable, or even enable the enhancement of desirable activities (16, 48). Therefore, a systematic means to identify functionally-relevant conformational transitions to excited states in the absence of stabilizing interactions could provide biophysical insight and new therapeutic opportunities. We expect the formation of a cryptic pocket to result in an exposon because, as explained above, the opening and closing of a pocket should result in cooperative changes in the solvent exposure of surrounding residues. Furthermore, for a cryptic allosteric site, we expect the allosteric coupling to give rise to correlations between the pocket exposon and residues around the relevant functional site.

We chose to test our approach on the enzyme TEM-1 β-lactamase because it is known to contain several cryptic allosteric sites (15, 17). It is also an important source of antibiotic resistance, so new inhibitors could provide a valuable means to restore the efficacy of existing antibiotics. In pursuit of new inhibitors, allosteric modulators have been discovered for at least one of these sites, which is created when a short alpha helix undocks from the protein, exposing a ligand binding site (15, 49, 50). To distinguish this site from other putative allosteric sites on this protein, we will refer to this site as the Horn pocket, or the Horn allosteric site, after the author who first reported this pocket (49).

As expected, we identify exposons corresponding to known cryptic pockets in TEM-1 β- lactamase (Fig. 2A). To visualize this, we mapped high total information exposons onto a crystal model of the TEM-1 ground state (Fig. 2B). In this format we observe a small number of spatially-condensed clusters of residues that are distant in sequence space, recapitulating our expectation that spatially (but not necessarily sequentially adjacent) objects are more likely to act cooperatively. The exposon with the highest total information (dark blue) corresponds to the active site. The exposon with the second-highest total information (light blue) corresponds to the Horn site (17, 49), shown in Fig. 2C, grey structure. Yet another exposon (beige, 4^th^ highest total information) reports on a second cryptic pocket that we reported previously (15). Each of the exposons corresponding to a cryptic pocket has substantial inter-exposon communication with the active site, suggesting the potential for perturbations to these pockets to exert allosteric control over activity.

**Figure 2:**
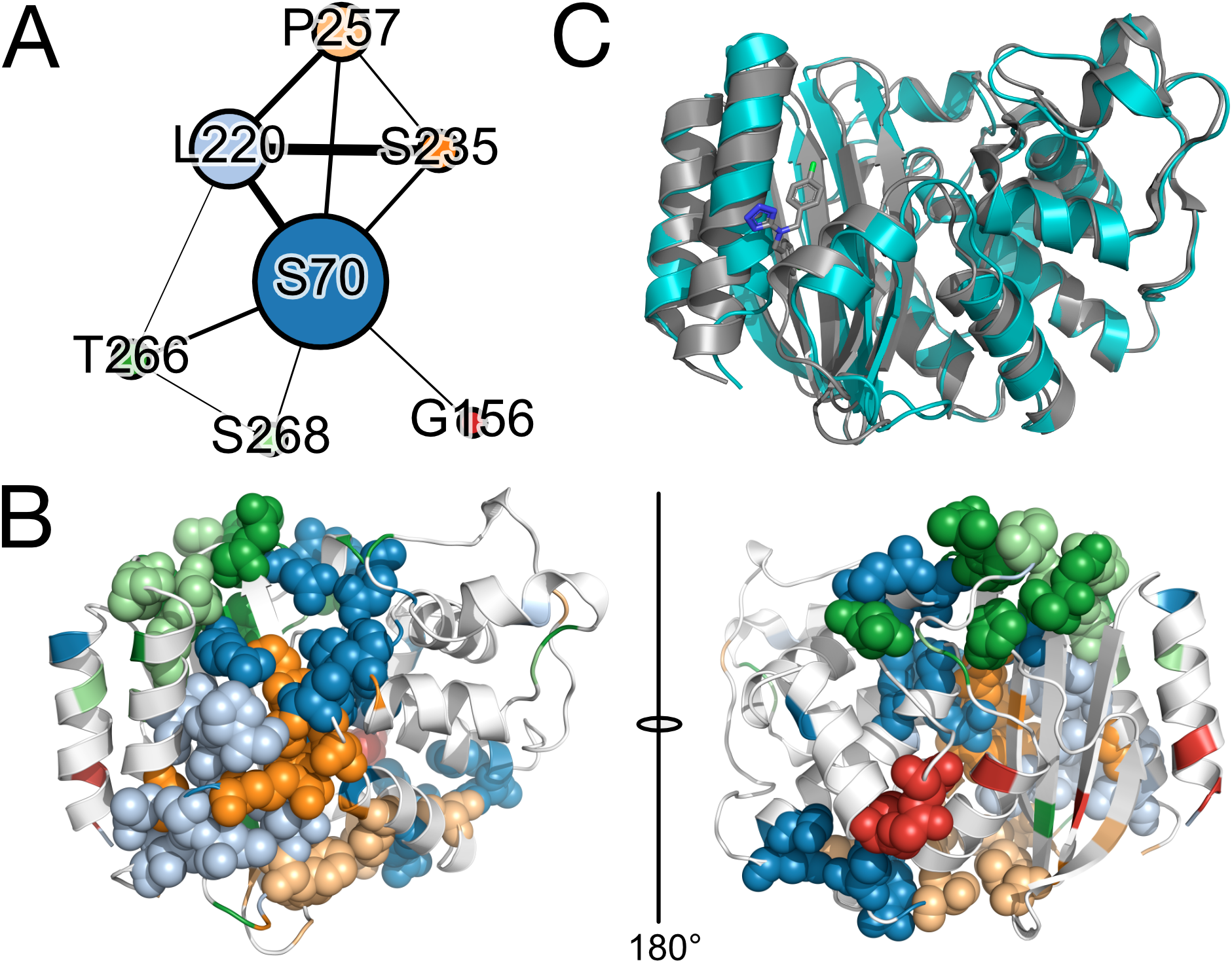
Exposons for TEM-1 β-lactamase. (A) The coarse-grained exposon network for TEM-1. Edge weights are proportional to the total correlation between each pair of exposons and node sizes are proportional to their eigenvector centrality. Self-edges and edges with very low values are omitted. (B) The highest total information exposons superimposed on a crystal model of unliganded TEM-1, 1JWP (64). Residue colors match the exposon colors in (A). Note the spatially contiguous exposons centered about the active site (dark blue), Horn allosteric site (light blue), a previously identified cryptic pocket (beige), and the Ω-loop (light and dark green). (C) A representative structure of the open state from the light blue exposon (teal) overlaid on a ligand-bound crystal structure of the Horn allosteric site (grey, 1PZO) (49).

To assess the effectiveness of using an MSMs’ eigenmodes to identify the motions that induce a particular exposon, we compare the structures identified in this way with known crystal models for the relevant ligand-bound state. In the case of the Horn cryptic allosteric site, a crystallographic model for the ligand-bound, open state is available (49). We then compare this structure with the structures at the extremes of the eigenmotion that best correlates with this exposon’s exposure state, as described in the Methods section. In this case, one extreme of the configuration resembles the ligand-free crystal structure, and the other is similar to the bound crystal structure (Fig. 1A, teal structure). The fact that the open structure from our model is somewhat more open than the ligand-bound structure is consistent with previous evidence that the pocket opens even further in solution than is seen in the crystal structure (17).

As an even more stringent test of our model, we assessed the consistency of our model’s predictions with an *in vitro* measurement of the kinetics of solvent exposure. Specifically, we used a thiol-labeling approach which we have improved from our previous work (15) by the use of a stopped-flow instrument (see *Methods*). In brief, this assay uses a drug-sized labeling reagent, DTNB (Ellman’s reagent) that changes absorbance upon covalently reacting with solvent-exposed reduced cysteines, providing a time-resolved measurement of residue-level solvent exposure with millisecond resolution. To make the comparison between our MSM and our thiol labeling experiment, we also developed a ‘synthetic labeling’ calculation (see *Methods*) that gives, as a function of time, the fraction of the population that has ever exposed the relevant sidechain to solvent.

As predicted, experimentally-confirmed pocket residues (S203, A232, L286) label at intermediate rates in our model (Fig. S7), while residues that remain buried in our experiments (L190, I260) do not show synthetic labeling and a surface control (A150) labels immediately in both our experiments and synthetic labeling. Rank is preserved: residues that label faster *in vitro* also label faster *in silico*. The main discrepancy between synthetic and experimental labeling occurs at S249, which labels very slowly *in vitro* but not *in silico*, likely because finite sampling prevented us from ever observing the slow process that leads to exposure of this residue. It is worth noting, however, this residue is located just “beneath” (i.e. deeper toward the core of the protein) an exposon that reports on cryptic pocket opening at this position (Fig. 2B, beige), suggesting that the exposon analysis may be somewhat robust to sampling error. In the future, a more precise understanding of the labeling reaction’s geometric requirements could enable quantitative predictions of labeling rates. For now, our model’s ability to correctly order pocket opening rates demonstrates its utility for identifying and characterizing pockets.

### Retrodiction of a conformational switch in nucleoprotein

As a subsequent test of our model, we assessed its ability to retrodict a conformational switch that was previously identified by Su et al (22). Proteins must frequently act as switches, altering their behavior in response to some signal. For example, many signaling proteins undergo conformational changes in response to specific stimuli that either increase or decrease their propensity to interact with downstream binding partners. We expect these concerted changes to manifest as exposons in our analysis.

As a test of the hypothesis that functional conformational switches at protein surfaces induce exposons, we analyzed Ebolavirus’s nucleoprotein (eNP), a conformational switch that controls access to and replication of the viral genome. Understanding and manipulating this conformational switch is of interest because Ebola was the causative agent in several recent, high case-fatality epidemics in sub-Saharan Africa (51) and is a pathogen for which very limited treatment options are available. Therefore, an improved biophysical understanding of this virus’s lifecycle may prove useful in understanding how to therapeutically target it. In one state, eNP oligomerizes and encapsidates the viral genome to package it for transport and protect it from degradation (52). In a second state, eNP exists as a monomer, releasing RNA to allow transcription of the viral genome (52). Recent evidence suggests that oligomerization is controlled by the curling of C-terminal helices of eNP into the RNA-binding cleft (22). We then expect an exposon to be formed by the residues in this groove and by the residues in the C-terminal tail that transiently occupies it.

Consistent with our expectation that the surficial rearrangements required for eNP function result in exposon formation, the highest total information exposon in eNP (Fig. 3A, dark blue) spans the residues in the C-terminal polymerization domain and the RNA binding groove. This is an interesting case in which an exposon would not be predicted to be composed of residues that are spatially contiguous when mapped to a crystal model of the ground state. This exposon is also at the center of the network of exposons (Fig. 3B). Extracting the motion that induces this exposon, shown in Fig. 3C, reveals that this exposon reports on the very same collective curling of the terminal helices into the RNA-binding cleft identified previously (53). Crucially, this dynamic process is consistent with hydrogen-deuterium exchange data that cannot be accounted for using available cryoelectron microscopy models (22). Manipulating this conformational equilibrium with small molecules or peptides could provide a powerful means of modulating the Ebola lifecycle. Indeed, a peptide that binds this interface has already been found to inhibit viral replication (53).

**Figure 3:**
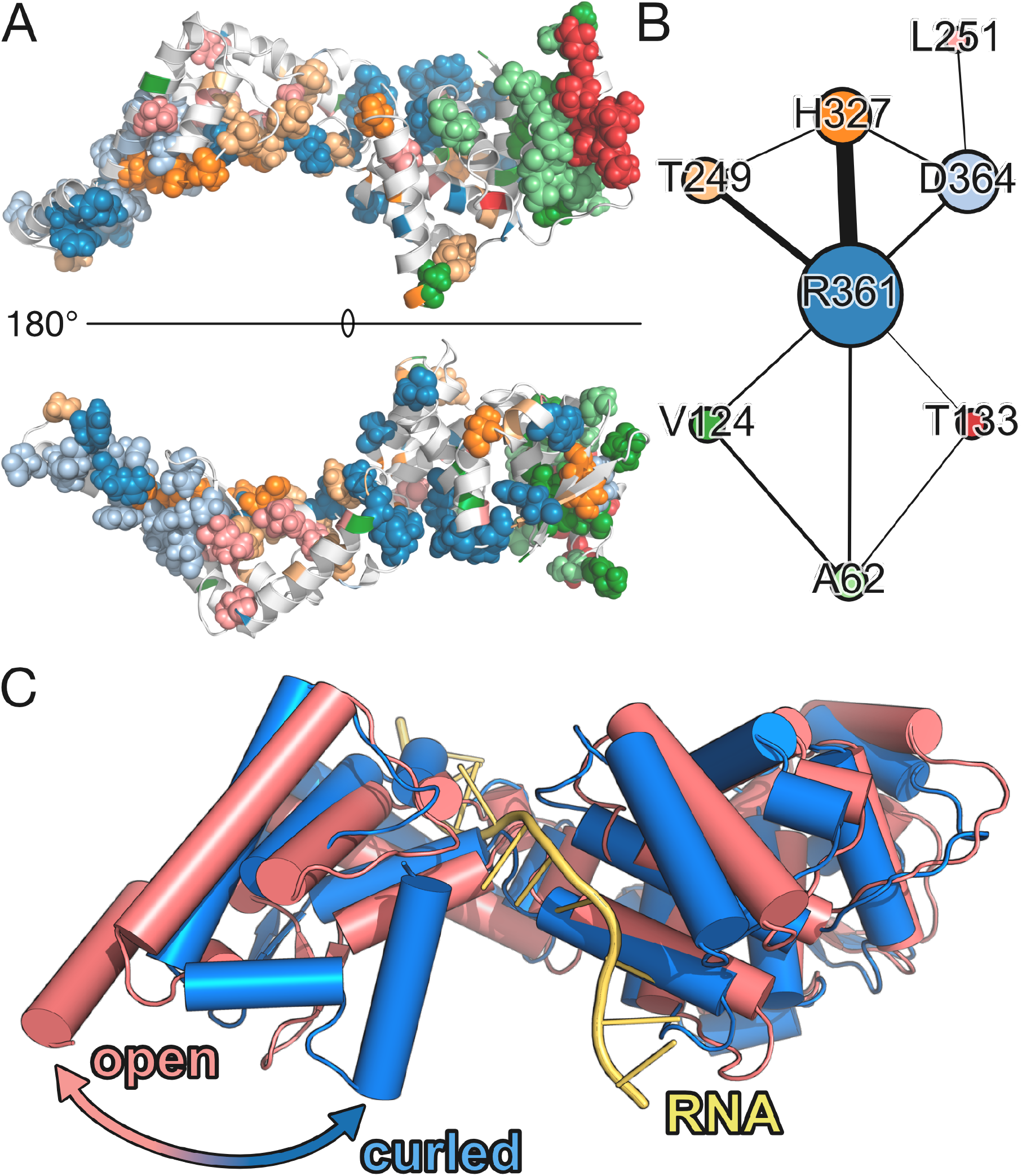
Exposons for eNP. (A) The distribution of the highest total information exposons superimposed upon a crystal model of monomeric eNP (22). Note the spatially non-contiguous nature of some of the exposons, especially the dark blue exposon. (B) The coarse-grained exposon network of eNP. Node colors match exposon colors in (A). Edge weights are proportional to the correlation between each pair of exposons and node sizes are proportional to their eigenvector centrality. Self-edges and edges with very low value are omitted. (C) the extremes of the eigenmotion best correlating with the highest total information exposon (blue exposon in A).

### Retrodiction of allosteric coupling between domains in CAP

As a further test of our model, we investigated its capacity to identify allosteric coupling between binding sites. Wherever an element of conformational selection is present, a binding site will sample both its bound and unbound configurations, and whenever the bound and unbound configurations differ in their pattern of solvent exposure, an exposon is expected to form. Because bound and unbound configurations presumably expose a different pattern of surface chemistry—one association-compatible and the other association-incompatible—we expect that differing patterns of solvent exposure might be a near-requirement. Furthermore, if these sites are allosterically coupled, they may even cluster into the same exposon.

Catabolite activator protein (CAP) is a homodimeric transcriptional activator in *E. coli* that allosterically couples cAMP binding to sequence-specific DNA association (54). This allosteric coupling between the cAMP binding domains (CBDs) and DNA binding domains (DBDs) is realized by a dramatic swiveling motion of the DBDs (55, 56), which changes the pattern of solvent accessibility on both the CBDs and DBDs, potentially producing one or more exposons. Besides coupling between the CBDs and DBDs (57), CAP also exhibits strong negative cooperativity between the two cAMP binding sites (58). Since these binding sites show different solvent exposure in cAMP-free and doubly cAMP-liganded crystal models, we expect these sites induce exposons as well. Since previous computational work on this protein suggests that evidence of this coupling is present in equilibrium simulations of the unliganded state (23), we expect to observe exposons that encompass residues in these regions.

As expected, the two highest total-information exposons computed from simulations of CAP in the unliganded state (Fig. 4A) are a symmetric pair stretching from the cAMP binding site in each monomer’s CBD to both DBDs. There is very strong communication between these two exposons, consistent with the negative cooperativity between the CBDs (54, 57).

**Figure 4:**
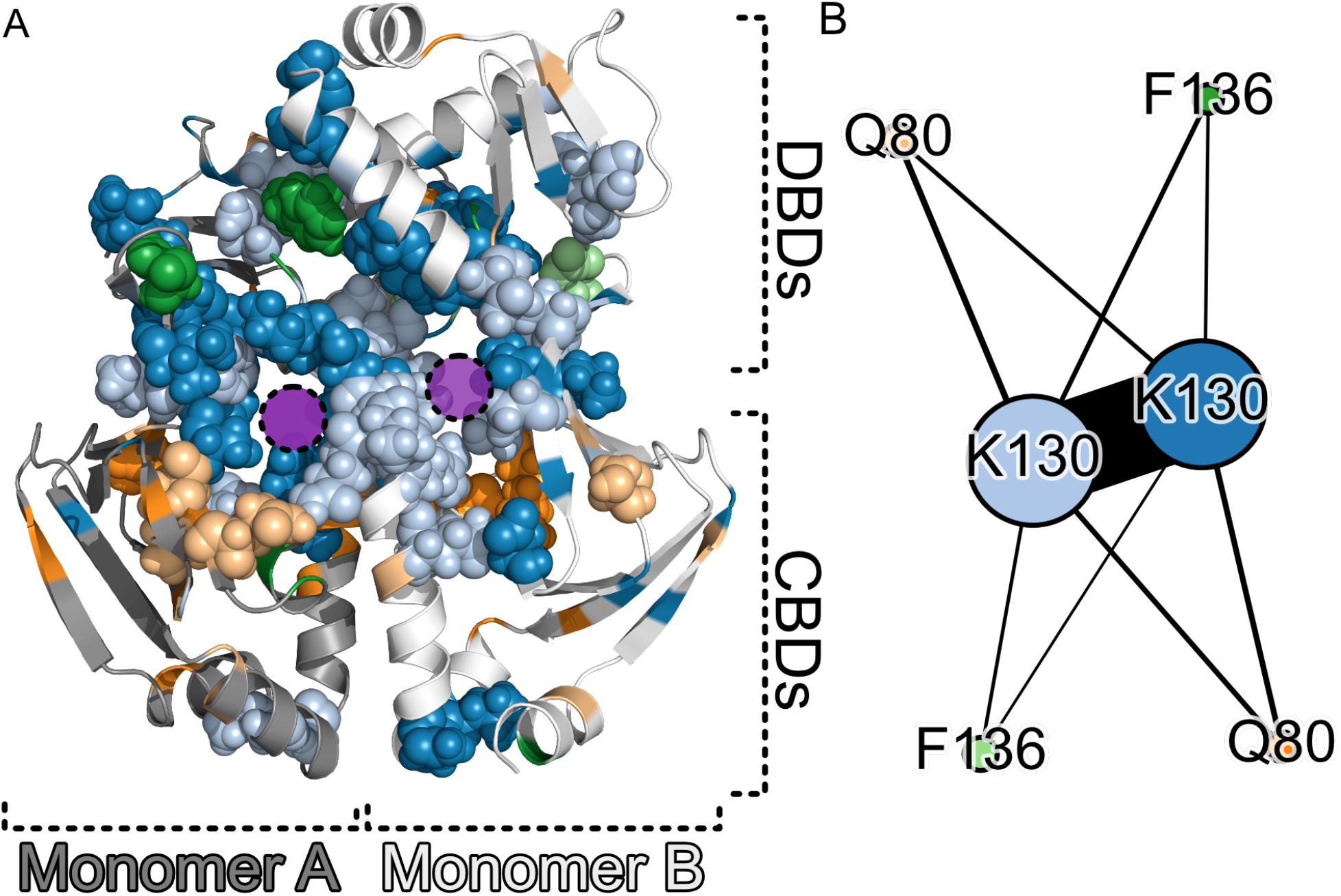
Exposons for CAP. (A) The distribution of the highest total information exposons superimposed upon a crystal model of unliganded, dimeric CAP (56). Purple circles indicate cAMP binding sites, and the cAMP-binding domains (CBDs) and DNA-binding domains (DBDs) are labeled. (B) The coarse-grained exposon network of CAP in graph form. Node colors match exposon colors in (A). Edge weights are proportional to the correlation between each pair of exposons and node sizes are proportional to their eigenvector centrality. Self-edges and edges with very low value are omitted.

The third-highest total information pair of exposons (Fig. 4A beige and orange) is centered about the individual cAMP binding sites, and they show less communication with one another than the larger DBD/CBD exposons (Fig. 4B). One explanation for the fact that these sites cluster separately from the rest of the cAMP binding site is that they are responsible primarily for substrate recognition, rather than allostery. The two highest total-information residues in this exposon, Q80 and R82, are two of the only four residues in the cAMP binding cassette that reduce their dynamicity upon binding (59)—opposite to the trend of the rest of the molecule and opposite to the hypothesized entropy-driven mechanism of allostery in this system. Furthermore, R82 is predicted to form a salt bridge with the cAMP phosphate and its mutation strongly affects binding (60).

To understand the motions that create the larger exposons reporting on interdomain and intermolecular allostery in this system, we examined the eigenmotion that best correlates with the highest total-information exposon we identified in this system (dark blue in Fig. 4A). A morph between the two extreme states of this eigenmotion (Video S1) indicates that this exposon represents a see-saw motion of the DBDs coupled to the closing of one cAMP site and the opening of the other. This is consistent with structural evidence (55) that the coupling between CBD and DBD involves large, rigid-body displacements of the two DBDs. This immediately suggests a testable hypothesis for how the negative coupling between cAMP binding sites might be achieved. This hypothesis could be further refined and dissected using methods like CARDS, as we have done previously (23), or experimental methods.

### Functional sites are exposon graph hubs

Exposons are a network model and consequently provide facile access to a protein’s allosteric topology. Because we have segmented the sequence into disjoint sets, this allows us to coarse grain our original mutual information matrix—which represents the sparse communication graph between all residues—into a much smaller graph representing communication between exposons. To calculate the communication between two exposons, we simply sum all edges that begin in one exposon and end in the other, and normalize by the channel capacity (23). The channel capacity is a measure of the maximum information that could possibly be transmitted between exposons, given the number of nodes they each contain (see *Methods*). Normalizing by this quantity allows for an intuitive comparison between the strength of communication between different pairs of exposons.

All exposon networks we examined had a hub-and-spoke architecture, with the exposon(s) with the highest total information serving as a hub and having a clear functional role. In TEM-1, the active site exposon (colored dark blue in Fig. 2B), including the active site serine, is visually central to the exposon graph (Fig. 5A), and each other node has its strongest connection with this node. We formalize this intuition by calculating each exposon’s eigenvector centrality (Fig. 5A-C) (61). Eigenvector centrality is a measure of the amount of time a random walker would spend at a particular node if transitions between nodes were distributed according to edge weights. Hence, nodes with higher-weighted or more connections to other nodes have a higher eigenvector centrality. In this case, we also find two groups of exposons attached to the hub but that are relatively uncorrelated with each other. Interestingly, one is a set of exposons that are under and around the Ω-loop, which is a critical modulator of substrate specificity and activity. In eNP, we also found that the exposon with the highest total information is a hub (Fig. 5B). As discussed previously, this exposon captures the dramatic curling motion that has been proposed to mediate RNA binding (22). In CAP, we find that the two exposons with the highest total information both have high centrality (Fig. 5C). These exposons appear to couple the ligand and DNA-binding domains of that protein.

**Figure 5:**
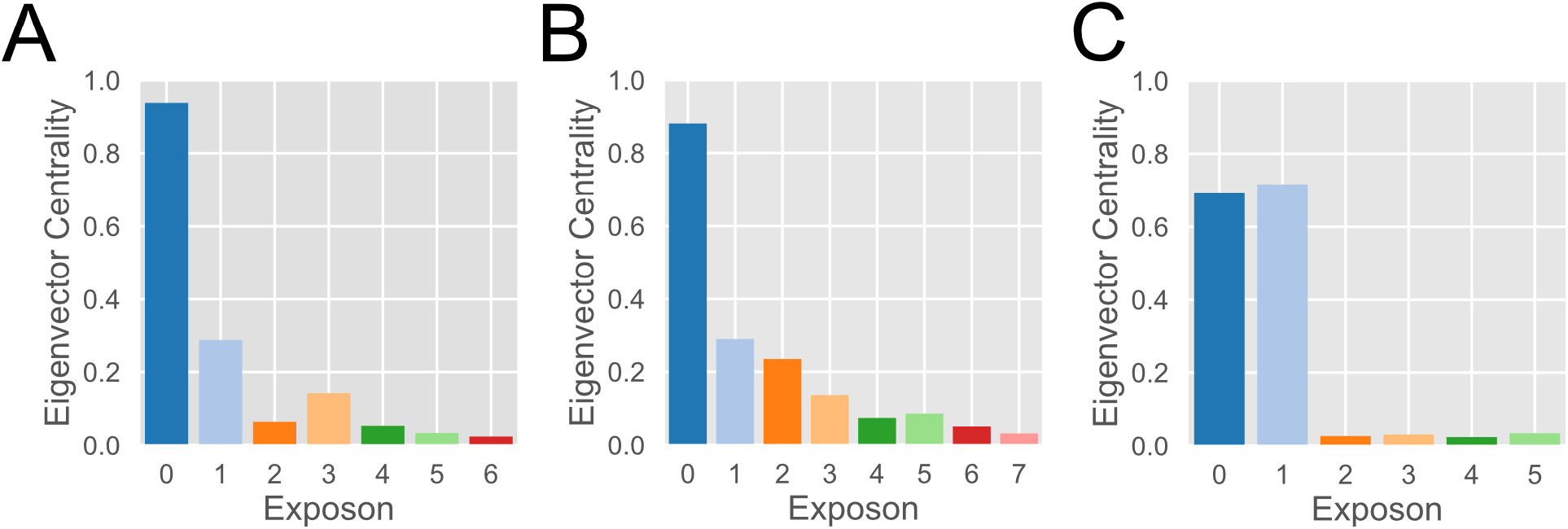
Eigenvector centrality of exposons for TEM-1, eNP, and CAP. Exposons are numbered from highest to lowest total information. In each case, the central exposon or exposons are associated with the primary function of the molecule they are found in. (A) In TEM-1, the most central exposon is at the active site. (B) In eNP, the central exposon is associated with a curling motion crucial to protein function (Fig. 3C). (C) In CAP, the pair of central exposons report on allosteric coupling between the DBDs and CBDs.

The fact that functionally-relevant conformational changes result in exposons with high total information and high centrality in three completely unrelated proteins furnishing wildly different functions is consistent with our motivating hypothesis that cooperativity does not arise at random.

### Discovery of the first known cryptic allosteric site in CTX-M-9

To demonstrate how the exposon model can be used to generate hypotheses and design experiments, we applied it to predict cryptic pockets in the enzyme CTX-M-9 β-lactamase. CTX-M-9 is interesting because, to the best of our knowledge, no cryptic pockets have been reported in this protein. It has less than 40% sequence identity with TEM-1, so it is not obvious whether or not it is likely to have similar cryptic pockets.

Examining the exposons for CTX-M-9 revealed that one of them contains the protein’s single native cysteine, C69 (Fig. 6A, yellow). This cysteine is completely buried in the apo crystal structure. Examining the motion that gives rise to this exposon reveals that C69 is exposed to solvent by a displacement of the Ω-loop (Fig. 6A), a structural element conserved among many β-lactamases and containing residues absolutely required for enzymatic activity (62) and that has significant conformational heterogeneity (37). The open structure of this pocket appears to be well-structured, as opposed to disordered, making it a potentially viable drug target. Therefore, we expect a small molecule that binds this pocket and displaces the Ω-loop would be a potent inhibitor while a drug that stabilizes the closed conformation would increase activity. The exposure of C69 in particular is of great interest because our thiol labeling assay can be applied without having to introduce a cysteine. Therefore, unlike previous applications of this method, there is no concern that the introduction of a cysteine created a pocket where none existed before.

**Figure 6:**
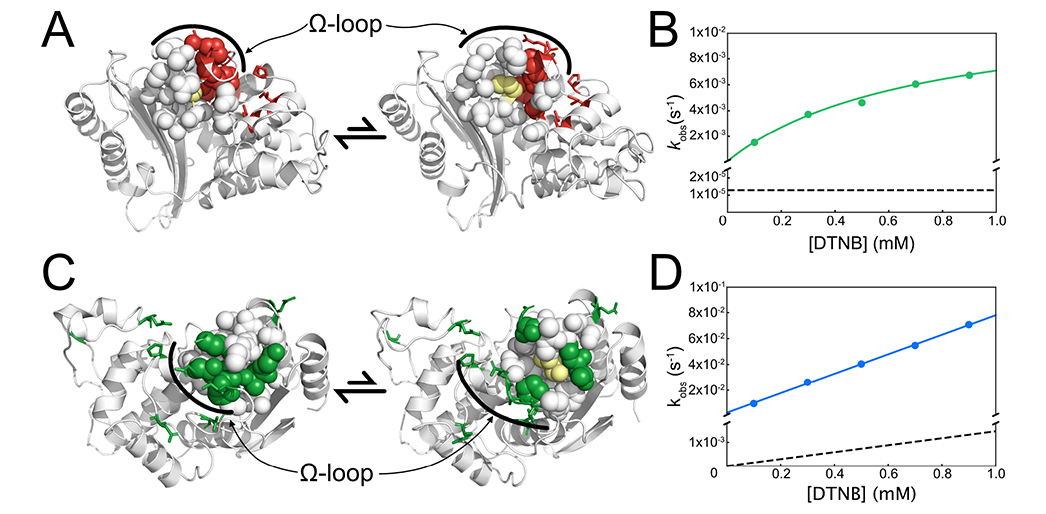
Exposons identify novel cryptic pockets in the CTX-M and TEM β-lactamases. (A) The extremes of the eigenmotion for the exposon containing C69 identify closed (left) and open (right) conformations of a cryptic pocket under the Ω-loop. Residues within 7 Å of C69 are shown as spheres and residues participating in the same exposon as C69 are shown as red sticks. Residue C69 is colored in yellow. (B), The observed labeling rates (solid green line) are in the EXX regime. The labeling rates expected for the global unfolding process (dashed line) are much slower. (C), the extremes of the eigenmotion best correlating with the exposon containing S243 identifies closed (left) and open (right) conformations of a cryptic pocket under and behind the Ω-loop in TEM-1. Residues within 7 Å of S243 are shown in spheres and residues participating in the same exposon as S243 are colored in dark green and as sticks. Residue S243 is colored in yellow. (D), The observed labeling rates (solid blue line) of a cysteine introduced at position 243 are in the EX1 regime. The labeling rates expected for the global unfolding process (dashed line) are much slower. In (B) and (D), standard deviations across three experiments were on the order of 10^-5^ and 10^-4^, respectively, and are not included for visual clarity.

We examined the labeling of C69 using our thiol labeling assay. The single exponential labeling that we observe is consistent with our prediction that C69 lines the first cryptic pocket to be identified in CTX-M-9 (Fig. 6B). C69’s labeling rate is much faster than the rate of the global unfolding process (Fig. 6B, dashed lines) measured by circular dichroism (Fig. S5), supporting our prediction that it is exposed by a fluctuation within the native state.

To ensure that exposon participation is a *bona fide* signal of pocket formation, we assayed the labeling rate of a residue that is buried in the crystal model but does not participate in an exposon, S123. Therefore, according to our model, a cysteine at this position should not show labeling. Consistent with this prediction, the S123C variant of CTX-M-9 does not show significant labeling.

To assess the allosteric potency of this site, we also measure the catalytic efficiency of the label-conjugated enzyme. In this case, after incubating CTX-M-9 with DTNB, which TNB-labels C69, we measure the rate at which it degrades nitrocefin, a β-lactam substrate (Fig. S7). TNB conjugation acts as a proxy for the binding of a drug. However, owing of TNB’s small size and hydrophilicity, this assay could easily underestimate the effect a true drug could have. We found an approximately 15-fold reduction in the catalytic efficiency (Fig. S7). By comparison, this same assay applied to previously-identified cryptic pockets in TEM-1 showed a less than threefold change in activity (15), making our newly identified site the most potent site in either TEM-1 or CTX-M-9.

Taken together, this newly-predicted pocket is the most attractive cryptic drug target site found to date in either TEM-1 or CTX-M-9. The fact that no mutation was required to perform thiol labeling of C69 also makes the results presented here some of the most compelling support for the predictive power of exposons in particular and MSMs in general.

### Prediction of a novel cryptic allosteric site in TEM-1

In light of our results for CTX-M-9, we examined the exposon graph for TEM-1 to see if a similar cryptic pocket may arise due to a displacement of the Ω-loop. Since TEM-1 has been extensively studied for the purpose of identifying cryptic allosteric sites, discovery of a new pocket in this molecule would be strong evidence for the utility of exposons for pocket discovery.

Two exposons (Fig. 2B, dark and light green), showing strong communication with one another, map onto the Ω-loop. The best-correlating MSM eigenmode revealed that S243 is significantly exposed by the opening of this pocket (Fig. 6C), and that the open form also appears well-structured and druggable. This conclusion is supported by quantitative druggability scores from fpocket (63) (Fig. S8). Interestingly, in our previous work, we were unable to detect this pocket because it frequently forms a channel-like connection with the active site, causing it to be combined with the active site pocket by pocket clustering methods (15, 17).

As position 243’s participation in an exposon predicts, the S243C variant labels at an intermediate rate that is slower than the near-instant labeling of a surface residue but substantially faster than the global unfolding process (Fig. 6D), which is on the order of hours (15). Once again, we also measured the catalytic function of the TNB-labeled enzyme.

Somewhat surprisingly, the TNB adduct had a 3.75-fold increase in catalytic efficiency—the ratio of k_cat_ over K_m_—driven primarily by a ∼4-fold decrease in K_m_ (Fig. S6, Table S3). This is consistent with recent evidence suggesting that both activation and inhibition are possible at the same allosteric site (50), and suggests that TNB may pack into the Ω-loop in such a way as to stabilize the closed conformation. Examination of crystal models of the closed state (64) reveals a void under the Ω-loop into which TNB might plausibly pack.

The fact that exposons identify a new cryptic allosteric site even in TEM-1—a protein that has been studied for many years by many groups, including intensively by our group with the specific goal of locating these sites—highlights the value of our approach for identifying functionally-relevant conformational changes. It also supports the hypothesis that the paucity of known cryptic allosteric pockets may stem more from technical limitations in locating them than from a low prevalence.

## Conclusion

We have demonstrated that exposons provide a powerful conceptual framework for identifying functionally-relevant conformational transitions. Exposons retrodict cryptic pockets, retrospectively identify conformational switches, and identify allosteric coupling between domains. We also showed that exposons are able to make *bona fide* predictions by discovering two new cryptic allosteric sites and experimentally verifying their existence. One of these sites is in a protein, CTX-M β-lactamase, that was not known to have any cryptic pockets, and in which no mutations were required to experimentally test our prediction. The other is in an enzyme that has been the target of an extensive search for cryptic pockets, so discovering a new site is a surprising testament to the power of exposons. Taken together, these results are compelling evidence for the utility of exposons.

Because many proteins’ most biologically interesting behavior involves changes at their surfaces, we expect our methodology to serve as a powerful first step in the analysis pipeline for proteins with complex, allosteric functions. Our results applying exposons to cryptic pockets, for example, demonstrates this method’s potential as the first step of a drug development pipeline targeting cryptic sites. Since the motions giving rise to exposons are substantially more diverse than simply pocket formation, exposons may also serve as a nearly automatic, high-throughput mechanism for dissecting allostery at protein surfaces either to refine an existing hypothesis or to identify potential alternative hypotheses.

Finally, the apparent ubiquity of the centrality of important functional surfaces in informational graphs suggests a general physical or biological principle in the organization of proteins. It was not obvious in our study exactly how this effect arises—reasonable hypotheses include genetic drift destroying any cooperativity that lacks function, or the thermodynamic cost of allosteric coupling when irrelevant—but it is universal amongst the proteins we have examined to date.

## Acknowledgements

We are grateful to the Folding@home users for computing resources. This work was funded by National Institutes of Health Grants R01GM12400701, U19AI109664, and T32GM02700, as well as by the National Science Foundation CAREER Award MCB-1552471. G.R.B. holds a Career Award at the Scientific Interface from the Burroughs Wellcome Fund and a Packard Fellowship for Science and Engineering from The David & Lucile Packard Foundation. M.I.Z. holds a Monsanto Graduate Fellowship and a Center for Biological Systems Engineering Fellowship.

